# Benchmarking Inverse Folding Models for Antibody CDR Sequence Design

**DOI:** 10.1101/2024.12.16.628614

**Authors:** Yifan Li, Yuxiang Lang, Chenrui Xu, Yi Zhou, Ziwei Pang, Per Jr. Greisen

## Abstract

Antibody-based therapies are at the forefront of modern medicine, addressing diverse challenges across oncology, autoimmune diseases, infectious diseases, and beyond. The ability to design antibodies with enhanced functionality and specificity is critical for advancing next-generation therapeutics. Recent advances in artificial intelligence (AI) have propelled the field of antibody engineering, particularly through inverse folding models for Complementarity-Determining Region (CDR) sequence design. These models aim to generate novel antibody sequences that fold into desired structures with high antigen-binding affinity. However, current evaluation metrics, such as amino acid recovery rates, are limited in their ability to assess the structural and functional accuracy of designed sequences.

This study benchmarks state-of-the-art inverse folding models—ProteinMPNN, ESM-IF, LM-Design, and AntiFold—using comprehensive datasets and alternative evaluation metrics like sequence similarity. By systematically analyzing recovery rates, mutation prediction capabilities, and amino acid composition biases, we identify strengths and limitations across models. AntiFold exhibits superior performance in Fab antibody design, particularly in variable regions like CDRH3, whereas LM-Design demonstrates adaptability across diverse antibody types, including VHH antibodies. In contrast, models trained on general protein datasets (e.g., ProteinMPNN and ESM-IF) struggle with antibody-specific nuances. Key insights include the models’ varying reliance on antigen structure and their distinct capabilities in capturing critical residues for antigen binding.

Our findings highlight the need for enhanced training datasets, integration of functional data, and refined evaluation metrics to advance antibody design tools. By addressing these challenges, future models can unlock the full potential of AI-driven antibody engineering, paving the way for innovative therapeutic applications.

**Author Summary:** Antibodies play a vital role in modern medicine, offering targeted therapies for diseases ranging from cancer to infectious diseases. Designing new antibodies with specific and enhanced functionalities remains a key challenge in advancing therapeutic applications. In this study, we benchmarked cutting-edge artificial intelligence models for antibody sequence design, focusing on their ability to generate sequences for the critical antigen-binding regions of antibodies, known as Complementarity-Determining Regions (CDRs).

Our findings reveal that specialized models like AntiFold excel in designing human antibody fragments, particularly in complex regions, while other models such as LM-Design demonstrate versatility across different antibody types. Importantly, we identified the limitations of models trained on general protein datasets, highlighting the need for antibody-specific training data to capture the unique features critical for therapeutic effectiveness.

By evaluating these models against robust datasets and diverse metrics, our work underscores the importance of improving training data and evaluation methods to advance AI-driven antibody design. These insights pave the way for more accurate and effective tools, ultimately supporting the development of next-generation antibody-based therapeutics.

## Introduction

Antibody-based therapies have revolutionized modern medicine, offering effective treatments for diseases ranging from cancer (e.g., pembrolizumab [1]), autoimmune disorders (e.g., infliximab [2]), hemophilia A (e.g., emicizumab [3] or mim8 [4]), and infectious diseases (e.g., palivizumab [5]). As demand grows for antibodies with enhanced functionality and tailored specificity, the ability to design novel antibodies rapidly and possessing the desired properties is increasingly critical for advancing next-generation biologics. Artificial intelligence (AI) holds tremendous potential to accelerate this process, providing innovative tools to tackle the challenges of antibody engineering.

Among these tools, antibody sequence design models, also known as “inverse folding” methods, have demonstrated significant progress in generating novel sequences that fold into desired structures with high antigen-binding affinity [6][7][8][9]. Historically, these models have been evaluated using amino acid recovery rates, which measure how accurately a model reproduces the exact native sequences of Complementarity-Determining Regions (CDRs)— key regions for antigen binding. While recovery rates provide a straightforward benchmark, they have notable limitations. State-of-the-art (SOTA) models have demonstrated remarkable progress, achieving amino acid recovery rates exceeding 50% for the Complementarity-Determining Regions (CDRs)—key regions for antigen recognition and binding [8][9][10].

However, amino acid recovery rates alone provide an incomplete assessment of model performance. First, recovery metrics penalize predictions that closely resemble native residues but differ by minor substitutions, such as lysine (K) to arginine (R), even though such substitutions often preserve structural and functional properties. Secondly, recovery metrics fail to account for the strong amino acid composition biases in CDR regions, such as the overrepresentation of glycine (G), serine (S), and tyrosine (Y). Models may exploit these biases to achieve high recovery rates without capturing the structural or functional constraints critical to antigen recognition. Finally, high-affinity binding often relies on a subset of critical residues within the CDRs, making it more relevant to evaluate a model’s ability to recover these key residues rather than all residues indiscriminately.

To overcome these limitations, we adopt sequence similarity as an alternative evaluation metric. Sequence similarity accounts for physicochemical properties, such as charge and hydrophobicity, allowing a more nuanced assessment of how well a designed sequence preserves native-like functionality. For instance, substitutions like lysine (K) to arginine (R) maintain positive charge and binding potential, making sequence similarity more reflective of practical design requirements. Furthermore, this metric prioritizes the accurate recovery of residues critical for antigen-antibody interactions, which are pivotal for therapeutic applications.

Some studies have also evaluated inverse folding models by assessing their ability to predict changes in affinity resulting from mutations but from a single dataset. In this study, we address this limitation by systematically benchmarking the models using a comprehensive collection of deep mutational scanning datasets but also to address the robustness of the different algorithms for antibody design. There are multiple different algorithms for designing antibodies such as: ProteinMPNN [6], ESM Inverse Folding (ESM-IF) [7], LM-Design [8], AntiFold [9], AbMPNN [10], IgMPNN [11], IgDesign [11], and Pi-Fold [12]. In this work, we have chosen to focus and benchmark these ones: ProteinMPNN, ESM-IF, LM-Design, and AntiFold due to their public availability and widespread use within the research community. By highlighting the strengths and weaknesses of these algorithms, our study provides insights into their robustness and offers a foundation for future improvements in antibody engineering tools.

## Methods

### Review of Architectures and Training data for Different Models

Recent advances in antibody inverse folding have resulted in the development of several computational tools with distinct architectures and training strategies. Below, we summarize the most widely used models:

#### 1. ProteinMPNN

ProteinMPNN, developed by the Baker lab, is a versatile model applied to diverse protein design tasks such as mini-protein binders and stable enzymes. The model utilizes a message-passing neural network architecture and is trained on high-resolution protein structures from the Protein Data Bank (PDB).

#### 2. ESM Inverse Folding

ESM Inverse Folding (ESM-IF) leverages a transformer architecture with Geometric Vector Perceptron (GVP) layers. It combines experimentally solved and predicted structures for training. By utilizing AlphaFold2, the authors expanded the training dataset to include predicted structures for approximately 12 million protein sequences, significantly increasing data diversity.

#### 3. LM-Design

LM-Design integrates sequence-based protein language models (pLMs) with structural encoders for sequence generation. It enhances the ESM-1b model with a lightweight structural adapter, incorporating structural awareness via ProteinMPNN. Unlike ESM-IF, LM-Design is trained solely on PDB-derived data, avoiding predicted structures.

#### 4. AbMPNN/AntiFold

AbMPNN and AntiFold are specialized models designed for antibody-specific inverse folding. Both models utilize a combination of experimentally solved structures from the Structural Antibody Database (SAbDab) and modeled antibody structures generated using ABodyBuilder2 from the Observed Antibody Space (OAS) paired database. The primary distinction between these models lies in their fine-tuning strategies: AbMPNN is fine-tuned on ProteinMPNN, whereas AntiFold is fine-tuned on ESM-IF. Our benchmarking primarily focused on AntiFold, as its authors reported superior performance compared to AbMPNN. Consistent with their findings, our internal data also indicates that AntiFold outperforms AbMPNN for Fab design. Detailed performance comparisons are available in Supplementary Figures 1 and 2.

### Evaluation Metrics and Tasks for Antibody Inverse Folding Models

To assess the performance of inverse folding models, we employed two key benchmarking tasks:

1. Antibody Sequence Design Each model was tasked with designing CDR sequences for given antibody-antigen complex structures. We use Fab to refer to two chain antibody binding fragments across the whole manuscript. For Fab antibodies, all six CDRs, three from the heavy chain (CDRH1(H1), CDRH2(H2), CDRH3(H3)) and three from the light chain (CDRL1(L1), CDRL2(L2), CDRL3(L3)) were designed. For VHH antibodies, only the three heavy chain CDRs (H1-H3) were included. To ensure fairness, the models were not conditioned on existing antigen or antibody framework sequences, as some models lack this capability. Preliminary experiments demonstrated that conditioning on framework sequences had minimal effect on sequence recovery (see Supplementary Figure 12).
1. Mutation Effects on Antibody Affinity: Predictions and Correlations Single-site mutagenesis was performed for all CDR residues, and models were used to calculate log-likelihood scores for each mutated sequence. These scores were then correlated with experimentally measured changes in binding free energy (ΔΔG) to evaluate the models’ ability to predict mutational effects.

### Evaluation Metrics for Antibody Inverse Folding Models

To quantify the success of the sequence designs, we used two metrics: design identity and design similarity.

1. Design identity is calculated as the sum of all perfectly recovered residues divided by the total number of designed residues.
2. Design similarity accounts for the physicochemical similarities between amino acids. We used the BLOSUM62 [13] substitution matrix to quantify the similarity between designed and wild-type (WT) residues. Design similarity is calculated as the sum of BLOSUM62 scores for all designed residues, normalized by the sum of BLOSUM62 scores for the WT residues.

The equation for design similarity is as follows:

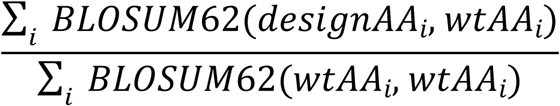

in which BLOSUM62(residueA, residueB) is the substitution score between resA and resB and i is the index of designed residues.

It is worth noting that some studies have utilized Rosetta energy scores to evaluate the quality of designs. However, we did not include this metric in our evaluation, as our preliminary analysis indicated only minor differences in energy scores between models. Additionally, the interactions present in the crystal structures themselves and the relaxation protocol appeared to have a greater influence on the energy terms.

### Antibody Structure Dataset and Preprocessing

We employed the Structural Antibody Database (SAbDab) [14,15] as the foundation for our dataset and applied rigorous filtering criteria to ensure data quality and relevance. These criteria included recency, sequence redundancy, structural quality, antigen relevance, and CDR3 uniqueness (see Table 1 for details). This filtering yielded 203 Fab and 61 VHH antibody structures (see Supplementary Tables 2 and 3 for PDB IDs and chains).

**Table 1:** Filtering criteria applied to the SAbDab dataset for antibody structure selection.

| Filtering Criteria | Description | Rationale |
| --- | --- | --- |
| Timeline Filtering | Included entries submitted after January 1, 2023 | Aligns with training data of the latest inverse folding models. |
| Sequence Redundancy | Excluded sequences with >95% identity to older entries | Reduces overfitting and ensures diversity. |
| Structural Quality | Removed structures with resolutions >3.5 Å or missing residues | Ensures high-quality templates for modeling. |
| Antigen Relevance | Excluded entries with multiple antigen chains or coronavirus antigens | Mitigates overrepresentation and simplifies modeling. |
| CDR3 Uniqueness | Retained unique CDR3 sequences; removed antibodies with identical CDR3 sequences. | Focuses on CDR3's pivotal role in antigen binding. |

### Sequence design Dataset

For our evaluation, we utilized the SAbDab dataset and applied rigorous filtering criteria to ensure data quality and relevance. These criteria are summarized in Table 2, detailing the specific steps taken to refine the dataset and their underlying rationale.

**Table 2:** Filtering criteria applied to the SAbDab dataset for sequence design evaluation.

| Filtering Criterion | Description | Rationale |
| --- | --- | --- |
| Temporal Filtering | Retained only entries submitted on or after January 1, 2023 | Focus on data relevant to recent advancements in antibody design. |
| CDR3 Sequence Filtering | Excluded sequences with CDR3 regions identical to those in pre-2023 PDBs | CDR3 is highly diverse and critical for antigen binding; redundant sequences reduce dataset value. |
| Duplication Removal | Remove duplicate sequences | Ensures a diverse dataset and avoids redundancy. |
| Antigen Filtering | Excluded entries lacking an antigen or with multiple interacting chains | Focuses on antibodies with well-defined antigen interactions. |
| Missing Residues | Removed structures with missing residues in antibody chains( e.g. those that cause broken chains in the crystal structure) | Prevents errors in sequence design models that rely on complete structures. |
| Resolution Criteria | Filtered out structures with a resolution higher than 3.5 Å | Ensures high structural quality for accurate modeling. |
| Similarity Filtering | Excluded PDBs with >95% sequence identity to pre-2023 SAbDab entries | Avoids overfitting and ensures the dataset reflects novel sequences. |
| Antigen Bias | Excluded PDB files with coronavirus antigens | Mitigates dataset bias, as ~30% of antibodies target coronaviruses. |

By applying these steps, we curated a high-quality dataset comprising diverse and recent antibody-antigen complexes suitable for benchmarking sequence design models. This filtering yielded 203 Fab and 61 VHH antibody structures (see Supplementary Tables 2 and 3 for PDB IDs and chains).

### Mutation Scoring Dataset

To benchmark the prediction of mutational effects, we curated datasets based on three criteria. First, the datasets must measure the impact of mutations on antibody-antigen binding, excluding any data related to antigen mutational scanning or mutations not associated with antibody binding complex. Second, each dataset must contain at least 20 mutation data points in the heavy chain to ensure that the calculated correlations have sufficient statistical power. Third, the dataset focuses only on heavy chain mutations with a fixed light chain, since a combination of heavy and light mutations will complicate the task. The datasets we collected include:

1. SKEMPI2 dataset [16]: contains mutation and relevant ddG. We could find 2 complex structures that have more than 20 mutation data points. We could calculate Per-Structure correlation of ddG and log likelihood for these pdb.
2. Affinity-matured influenza broadly neutralizing antibodies (bnAbs) CR9114 and CR6261 [17]: The profiled mutational landscape of CR9114 includes all possible combinations of 16 substitutions, whereas that of CR6261 includes all possible combinations of 11 substitutions, totaling 65,536 and 2048 variant antibody sequences, respectively.
3. Shira Warszawski performed saturation mutagenesis for all CDR residues for an anti-lysozyme antibody, and used yeast display and deep sequencing to estimate the affinity of each mutation [18].

### Antibody Sequence Design and Residue Classification Sequence Design Protocol

For sequence design, PDB files were preprocessed to retain only the Fv region (for Fab) or the VHH chain, along with the antigen chain. Complexes with multiple antigen chains were excluded. CDR regions, defined using the Chothia numbering scheme [19] with the AbNumber tool [20], were designed without conditioning on antigen or framework sequences due to limitations in ESM-IF and AntiFold. For antigen-free design, the antigen chain was removed from the input PDB file. For relaxed structure design, PDBs underwent three rounds of Rosetta FastRelax with coordinate constraints (weights = 1.0) [21–23](see Supplementary Method 1 for detail relaxation xml file). Sequence recovery and similarity were calculated for each CDR region and PDB file.

### Categorization of CDR Residues by Location and Functional Role

To investigate the relationship between residue location, function, and design accuracy, we categorized CDR residues based on their relative solvent accessible surface area (SASA) and predicted contribution to antigen binding.

### SASA Calculation and Buried Residues

SASA was calculated for each residue in the unbound antibody (apo) format using the Shrake-Rupley algorithm. Residues with a relative SASA below 20% were classified as “buried,” indicating limited solvent exposure.

### Predicting Binding Contributions and Key Interactions

We employed an Rotamer Density Estimator (RDE) based mutation effect prediction model to predict the contribution of each residue to the binding free energy (ΔΔG). This model was internally fine-tined (unpublished) with a network pretrained to predict the probability density of protein side-chains and a module to estimate the mutational ΔΔG with the representations learned from the pre-trained model. To estimate the contribution of each residue for antigen binding, we performed an in silico alanine scanning to all residues in the CDR regions. More specifically, we mutate the native residue to alanine for all CDR residues and calculated the ΔΔG for the mutation. A higher ΔΔG indicated a larger drop of binding affinity with the mutation. We define residues with a ΔΔG greater than 1.5 kcal/mol upon alanine substitution as “key interaction” residues, representing around 5% of all the residues in the CDR regions.

### Surface Contact Residues

Residues located within 4 Å of any antigen atom but not classified as key interaction residues were categorized as “surface contact” residues. These residues are solvent-exposed and in contact with the antigen but are not predicted to make major energetic contributions to binding.

This categorization scheme allowed us to assess the models’ performance in designing sequences for residues with distinct structural and functional roles.

## Results

### Benchmarking Antibody Design Models: Recovery Rates and Similarity Across CDRs

#### Performance Across Fab Antibodies

We evaluated the sequence recovery and similarity of CDRs designed by each model for Fab antibodies. The models ranked in performance as follows: AntiFold > LM-Design > ESM-IF ≈ ProteinMPNN (Figure 1A). Notably, AntiFold consistently outperformed the other models, achieving higher recovery rates and sequence similarity across all CDR regions except for H3 (see Supplementary Figure 1 and Supplementary Table 4 for detailed description of recovery rates). LM-Design demonstrated robust performance, particularly in recovering residues within canonical CDR regions, where it surpassed ESM-IF and ProteinMPNN. However, its recovery rates for H3 were slightly lower than AntiFold, which excelled in capturing the structural diversity of this region. These findings underscore AntiFold’s ability to design sequences that align closely with native structures, particularly in challenging regions like CDRH3.

**Figure 1:**
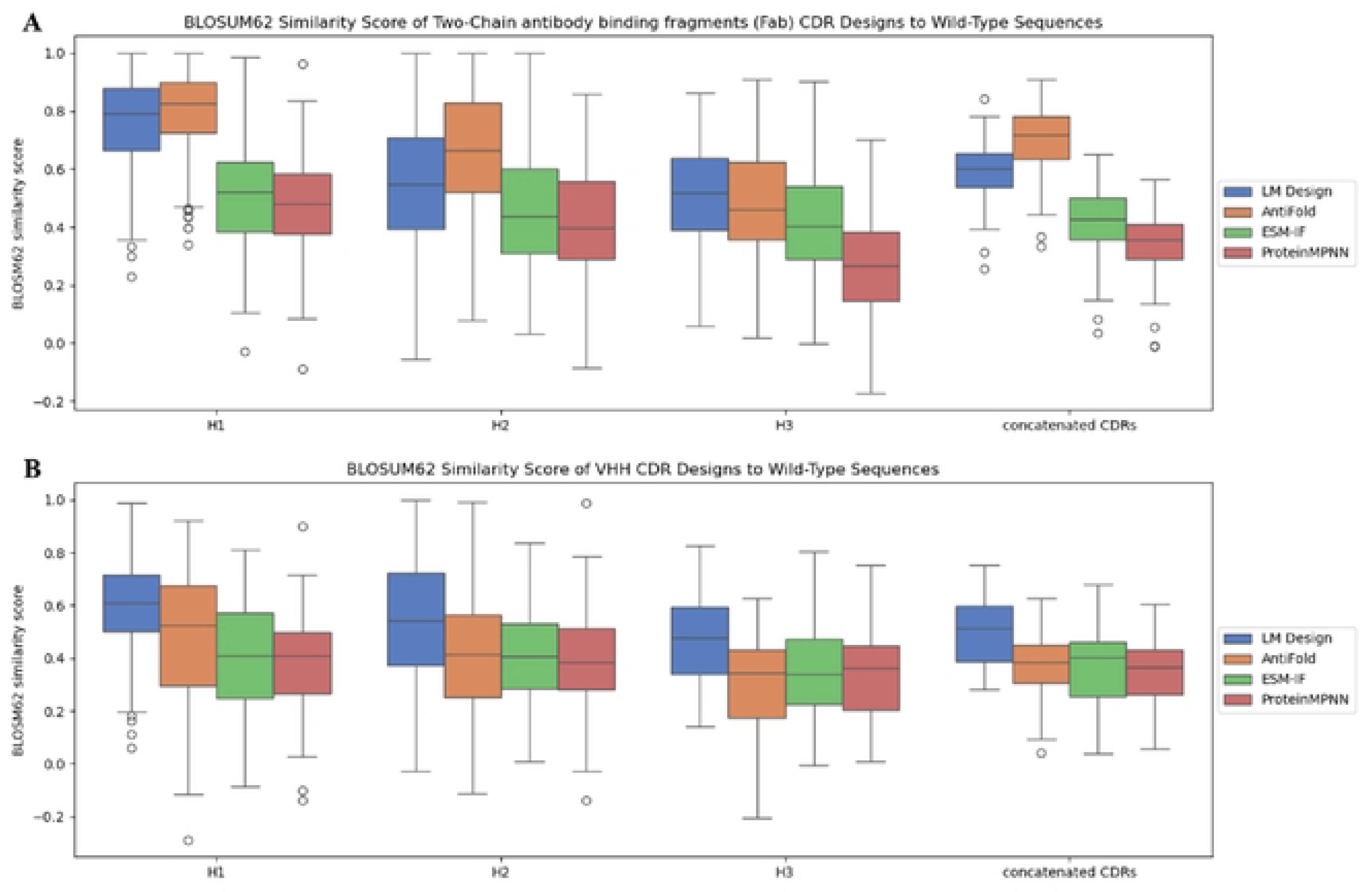
Design methods compared: LM Design, AntiFold, ESM-IF, and Protein MPNN, using BLOSUM62 similarity scores. Models were evaluated on 203 Fab (panel A) and 61 VHH (panel B) crystal structures from SAbDab. Analyses were restricted to H1, H2, and H3 regions, as defined by the Chothia numbering scheme.

#### Performance Across VHHs

For VHH antibodies, the performance ranking shifted: LM-Design > AntiFold ≈ ESM-IF ≈ ProteinMPNN (Figure 1B). LM-Design achieved the highest recovery and similarity scores across all CDR regions, highlighting its adaptability to the compact and simplified architecture of VHH antibodies. In contrast, AntiFold, which performed exceptionally well for Fab antibodies, showed reduced efficacy for VHH sequences. This discrepancy likely stems from its training data, which lacked substantial representation of VHH antibodies. Despite this limitation, AntiFold maintained competitive performance in H1 and H2, demonstrating some generalizability across antibody types (Supplementary Figure 2 and Supplementary Table 5).

#### CDR-Specific Trends

The recovery rates and similarity scores varied significantly across the six CDR regions in Fab antibodies and three in VHH antibodies. AntiFold showed remarkable performance in H1 and H2 for both Fab and VHH, reflecting its strength in designing residues within relatively conserved regions. However, its performance for H3 was mixed, excelling in Fab antibodies but lagging in VHH sequences.

LM-Design’s performance was more uniform across all CDRs, particularly excelling in the highly variable H3 of VHH antibodies. ESM-IF and ProteinMPNN showed comparable recovery rates, with both models performing better in light chain regions (L1, L2, L3) than in heavy chain regions. These trends are summarized in Figure 2, Supplementary Figure 1 and 2, and Supplementary Table 4 and 5.

**Figure 2.**
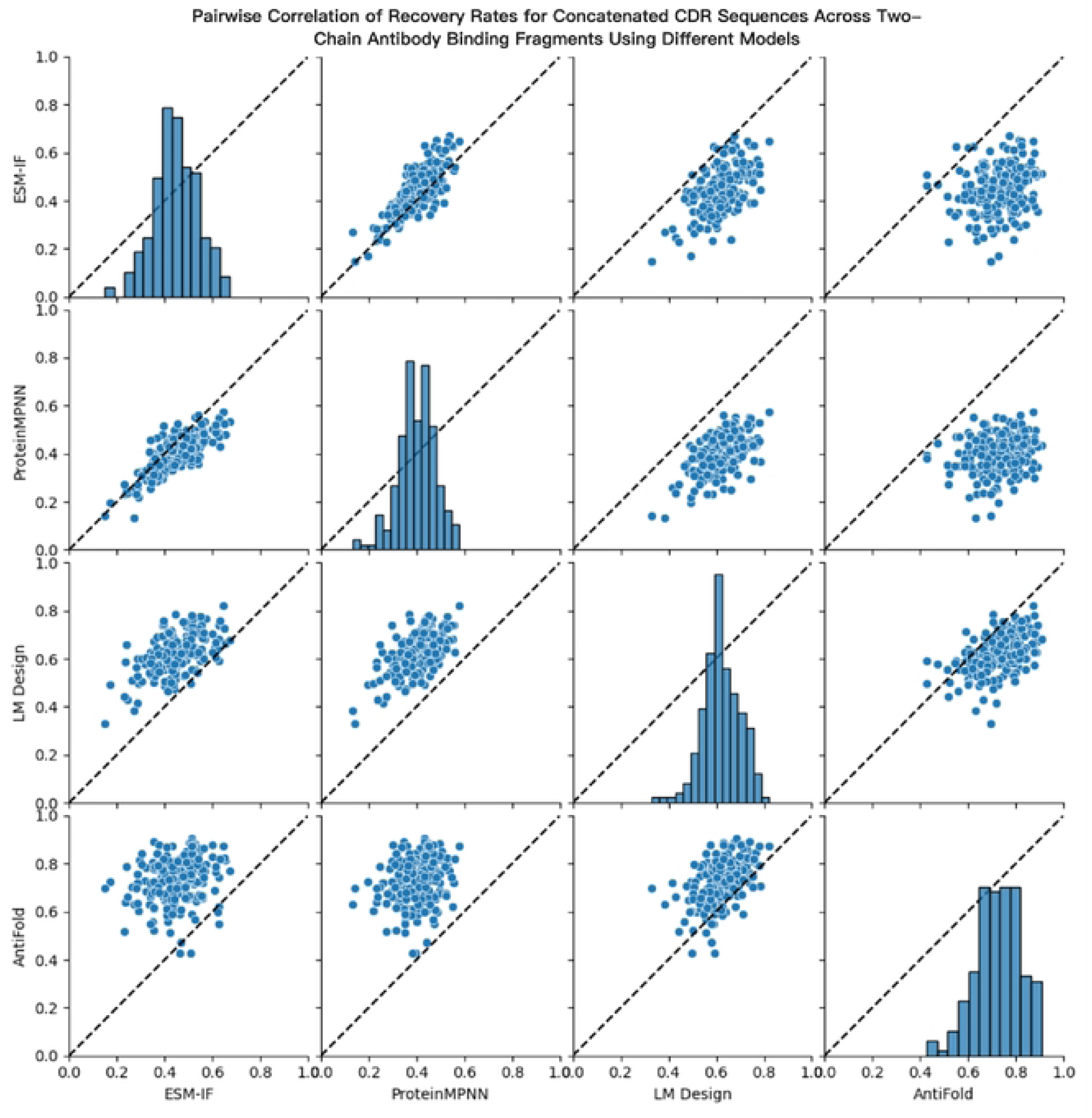
Pairwise correlation of recovery rates for concatenated CDR sequences across two-chain antibody binding fragment PDB structures using various models. Each dot represents the recovery score for a single PDB stn1cture in the benchmarking dataset. Recovery or similarity scores were calculated for all concatenated CDR regions.

#### Key Insights

##### AntiFold’s Unique Strengths

AntiFold’s fine-tuning on antibody-specific datasets contributed to its superior performance in Fab antibodies, particularly in capturing the variability of H3 (Supplementary Table 4). Its reduced reliance on antigen context further enhances its robustness for antigen-independent tasks.

##### LM-Design’s Adaptability

LM-Design’s integration of protein language models allowed it to generalize effectively across both Fab and VHH antibodies. Its balanced performance across CDRs highlights its potential for designing antibodies with diverse structures.

##### Challenges for General Protein Models

ESM-IF and ProteinMPNN, trained primarily on general protein datasets, struggled to capture the unique sequence-structure relationships of antibodies. These models exhibited biases toward overrepresented residues and failed to effectively design variable regions like H3.

These insights are summarized in Figure 2, Supplementary Figure 1 and 2, and Supplementary Table 4 and 5.

#### Analysis of Amino Acid Prediction Accuracy

To assess the models’ ability to recover specific amino acid types within the CDRs, we analyzed the prediction accuracy for each residue type. Overall, AntiFold demonstrated the highest recovery rates, followed by LM Design, and then ESM-IF and ProteinMPNN, which showed comparable performance (Figure 3 and Supplementary Figure 10). Notably, AntiFold outperformed LM Design in recovering almost every amino acid type, except proline and glycine.

**Figure 3.**
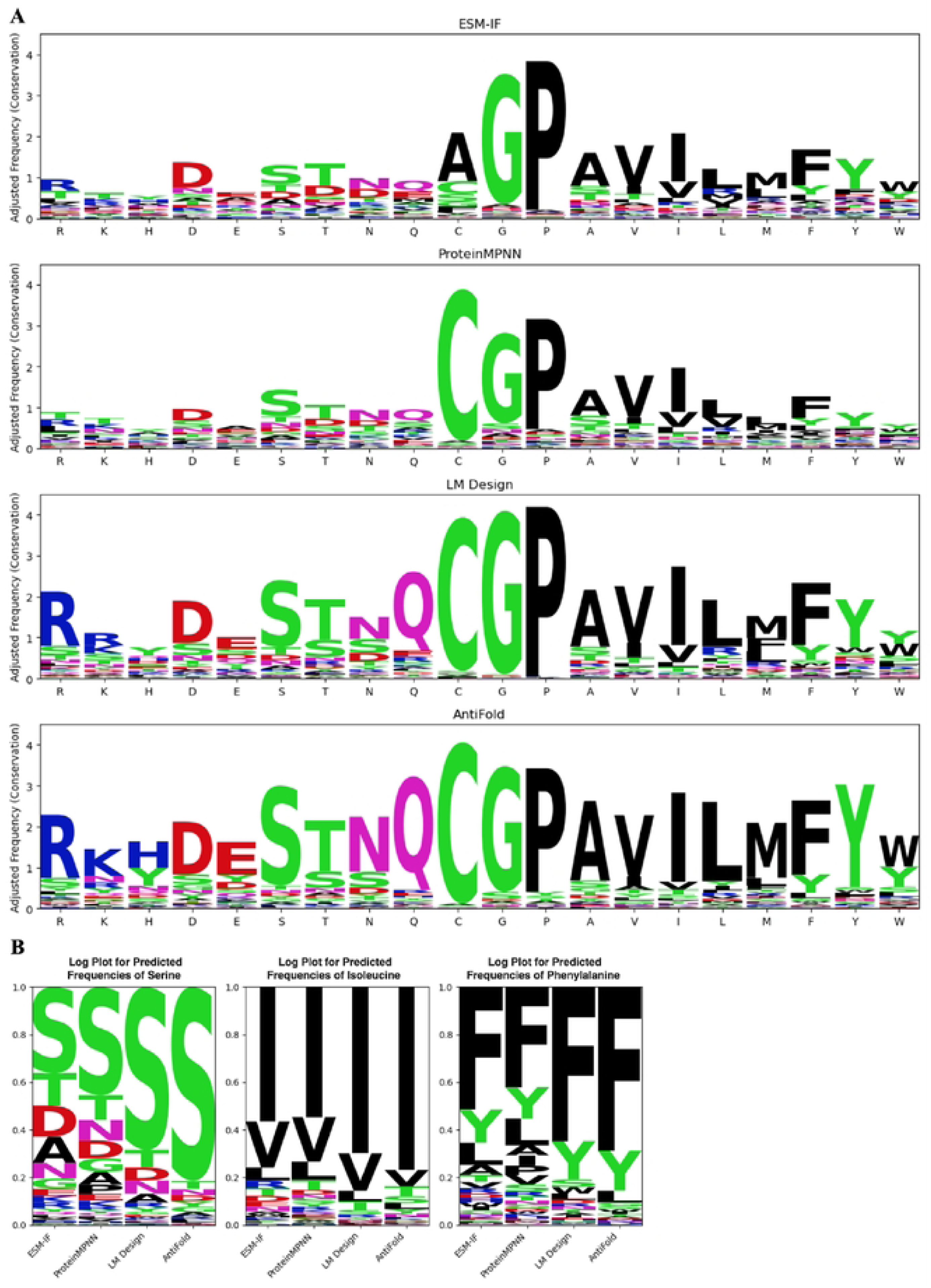
Analysis of Amino Acid Prediction Accuracy in Two-Chain Antibody Binding Fragment CDR Design A) Logo plots showing the predicted amino acid frequencies for each wild-type (WT) residue in the CDRs of two-chain antibody fragments. WT residues arc ordered by descending frequency. The height of each letter indicates the relative frequency of predicted amino acids, highlighting each model’s substitution tendencies. The overall height of each column indicates the conservativeness of the design for each type of residue. See Supplementary Figure 11 for a corresponding analysis of VHH CDR designs. B) Focused logo plots displaying prediction frequencies for three specific residue types (S, I, and F) across different models. Amino acid predictions are sorted from highest to lowest frequency for each residue type.

Further examination revealed an interesting trend across all models. Even when incorrect predictions occurred, the substituted residues often exhibited similar physicochemical properties to the WT residues. Common substitutions included those within the following groups: lysine (K) / arginine (R), threonine (T) / serine (S), and tryptophan (W) / phenylalanine (F) / tyrosine (Y). We selected three representative amino acid types—S, isoleucine(I), and F — and calculated the frequency of predicted types in Figure 3B. It is evident that for all models, the most frequent predictions correspond to the native residues, while the second most frequent predictions exhibit chemical properties closely resembling those of the native residue (e.g., Y for F, valine(V) for I, and T for S). This observation suggests that the models have learned to recognize and preserve key amino acid properties during sequence design, even if they do not always predict the exact native residue.

#### Analysis of Amino Acid Frequency Bias in Designed Sequences

To assess whether the models accurately capture the natural amino acid distribution within CDRs, we analyzed the frequency of each residue type in the designed sequences. For Fab antibody design, LM Design and AntiFold exhibited a stronger correlation with natural amino acid frequencies compared to ProteinMPNN and ESM-IF. This suggests that LM Design and AntiFold more effectively mirror the observed biases in amino acid usage. For example, both models accurately captured the overrepresentation of phenylalanine and serine in CDR regions, which was less pronounced in the sequences generated by ProteinMPNN and ESM-IF.

In contrast, for VHH antibody design, LM Design aligned closely with natural amino acid frequencies, whereas AntiFold deviated significantly. AntiFold exhibited a tendency to overrepresent high-frequency residue types in its designs (Figure 4 and Supplementary Data). This discrepancy suggests potential differences in how AntiFold models Fab and VHH sequences. We hypothesize that AntiFold, trained predominantly on a Fab dataset, may attempt to approximate human IGHV3 sequences when designing VHH formats, as IGHV3 is the closest match in its training data. Supporting this hypothesis, AntiFold-designed VHH sequences exhibited amino acid biases similar to IGHV3 sequences, including a higher frequency of Y, D, F, G, and S, and a lower frequency of R and T. For details, refer to Figure S5 and the findings presented in Gordon et al [24].

**Figure 4.**
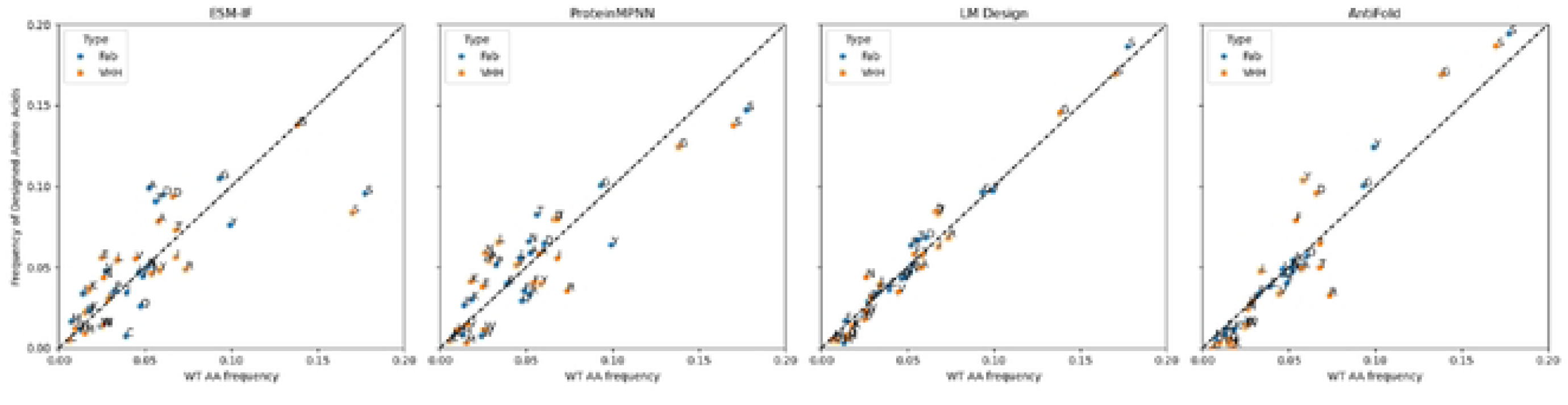
Amino Acid Frequency Comparison Between Designed and Wild-Type Sequences. Scatter plots comparing the amino acid frequencies of designed sequences to their wild-type (WT) counterparts for Two-Chain antibody binding fragments: Fab (blue circles) and VHH (orange circles). The frequencies were calculated by counting all WT and designed residue types across the benchmarking dataset. The diagonal dashed line indicates a perfect match between designed and WT frequencies.

**Figure 5.**
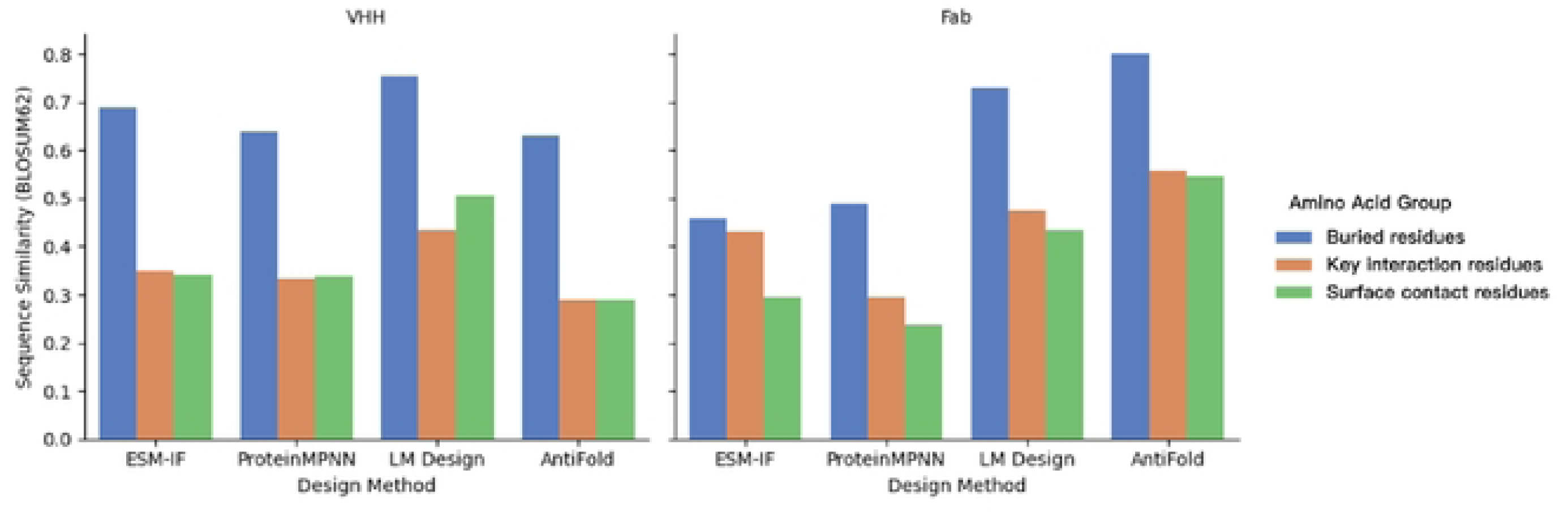
Sequence Design Similarity for Residues in Different Functional Regions of Two-Chain Antibody Binding Fragments (Fab, right) and Single-Domain Antibodies (VHH, left) The BLOSUM62 similarity scores were calculated for residues in three key regions: buried, key interaction, and surface contact. Residues were grouped based on their location and functional role in the complex. Sequence similarity was determined by comparing all residues in the respective regions of the designed sequences to their wild-type counterparts.

**Figure 6.**
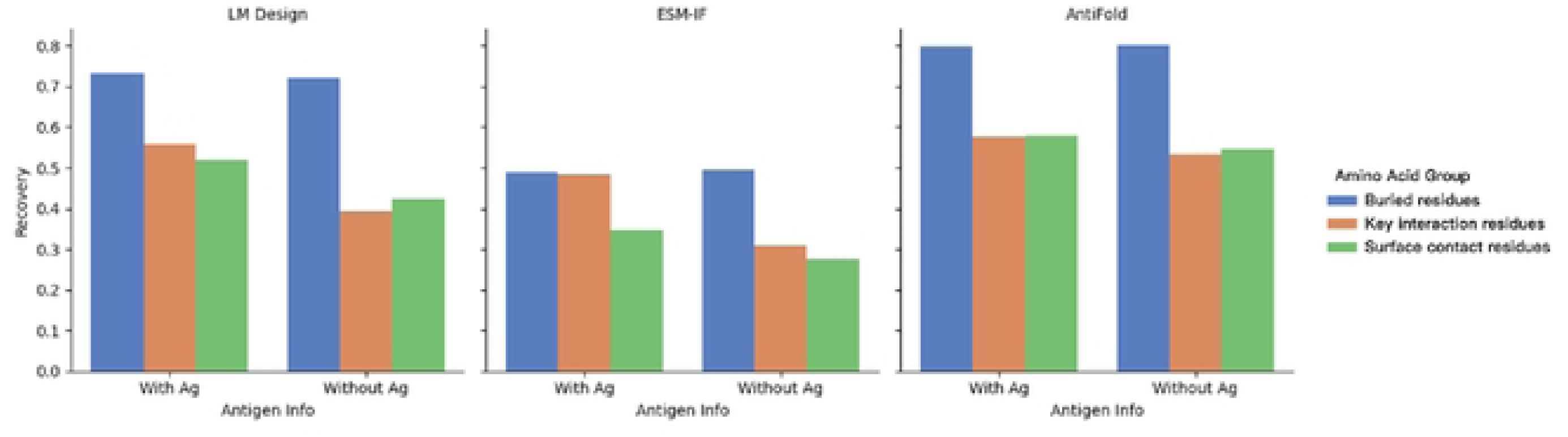
Sequence Design Recovery Rates With and Without Antigen Presence Recovery rates for designed sequences were analyzed under two conditions: with the antigen present in the structure (With Ag) and after the antigen chain was removed from the PDB file (Without Ag). Residues were grouped into three functional categories: bu1ied, key interaction, and surface contact. The bars indicate the recovery rates for residues in these regions across different models (LM Design, ESM­IF, and AntiFold), highlighting the impact of antigen presence on sequence design accuracy.

#### Sequence Design Performance by Residue Location and Role

Recognizing that amino acid residues in different regions of an antibody play distinct functional roles, we evaluated the models’ ability to design sequences based on residue location and role in antigen binding. We categorized CDR residues into three groups:

- Buried: Residues with minimal solvent exposure in the unbound antibody (apo) format.
- Key Interaction: Residues identified as critical for antigen binding based on their RDE score, a measure of the impact of alanine mutations on binding affinity.
- Surface Contact: Residues located on the antibody surface that contact the antigen but are not classified as key interaction residues.

Detailed criteria for these classifications are provided in the Methods section.

We then calculated the design similarity and recovery rates for each residue group. Across all models and antibody types, buried residues exhibited the highest design similarity, likely due to their greater conservation compared to surface residues. For Fab antibodies, AntiFold and LM Design showed significantly higher recovery rates for buried residues than ESM-IF and ProteinMPNN.

Interestingly, performance differences between models were less pronounced for key interaction residues. Notably, ESM-IF achieved similarity values for key interaction residues comparable to buried residues. Moreover, ESM-IF showed the largest difference in similarity between key interaction residues and surface contact residues. These findings suggest that ESM-IF may be particularly sensitive to the structural context of the antigen-binding interface.

#### Impact of Antigen Structure on Sequence Design

To investigate the influence of the antigen’s structural context on the models’ sequence design capabilities, we compared sequence recovery rates with and without the antigen chain present in the input PDB structure. This allowed us to assess the extent to which each model relies on the antigen’s presence for accurate sequence design.

We found that LM Design and ESM-IF produced slightly lower sequence design similarities when the antigen was excluded, indicating that these models utilize antigen information to some extent. In contrast, AntiFold showed nearly identical design similarities regardless of the antigen’s presence, suggesting that it primarily relies on the CDR loop structure for sequence design.

To further explore this observation, we analyzed the recovery rates for different groups of residues (buried, key interaction, and surface contact) with and without the antigen present. As expected, the recovery rates for buried residues remained consistent regardless of the antigen’s presence. However, for key interaction and surface contact residues, the models exhibited varying degrees of dependence on the antigen structure. ESM-IF showed a significant drop in recovery rates for these residues when the antigen was absent, highlighting its reliance on antigen information for designing interface residues. Conversely, AntiFold’s performance remained consistent, reinforcing the observation that it primarily designs sequences based on the antibody’s structure rather than its interaction with the antigen.

#### Illustrative Design Examples and Model-Specific Performance

To gain deeper insights into the performance of different models, we analyzed the logo plots of several representative designs. PDB 8TFL, which depicts a neutralizing Fab fragment SylH3 bound to ricin toxin, serves as an illustrative example. We found that AntiFold and LM Design achieved nearly 90% recovery rates for H1, whereas ProteinMPNN and ESM-IF exhibited recovery rates of only around 40% (Figure 7 and Supplementary Table 1).

**Figure 7.**
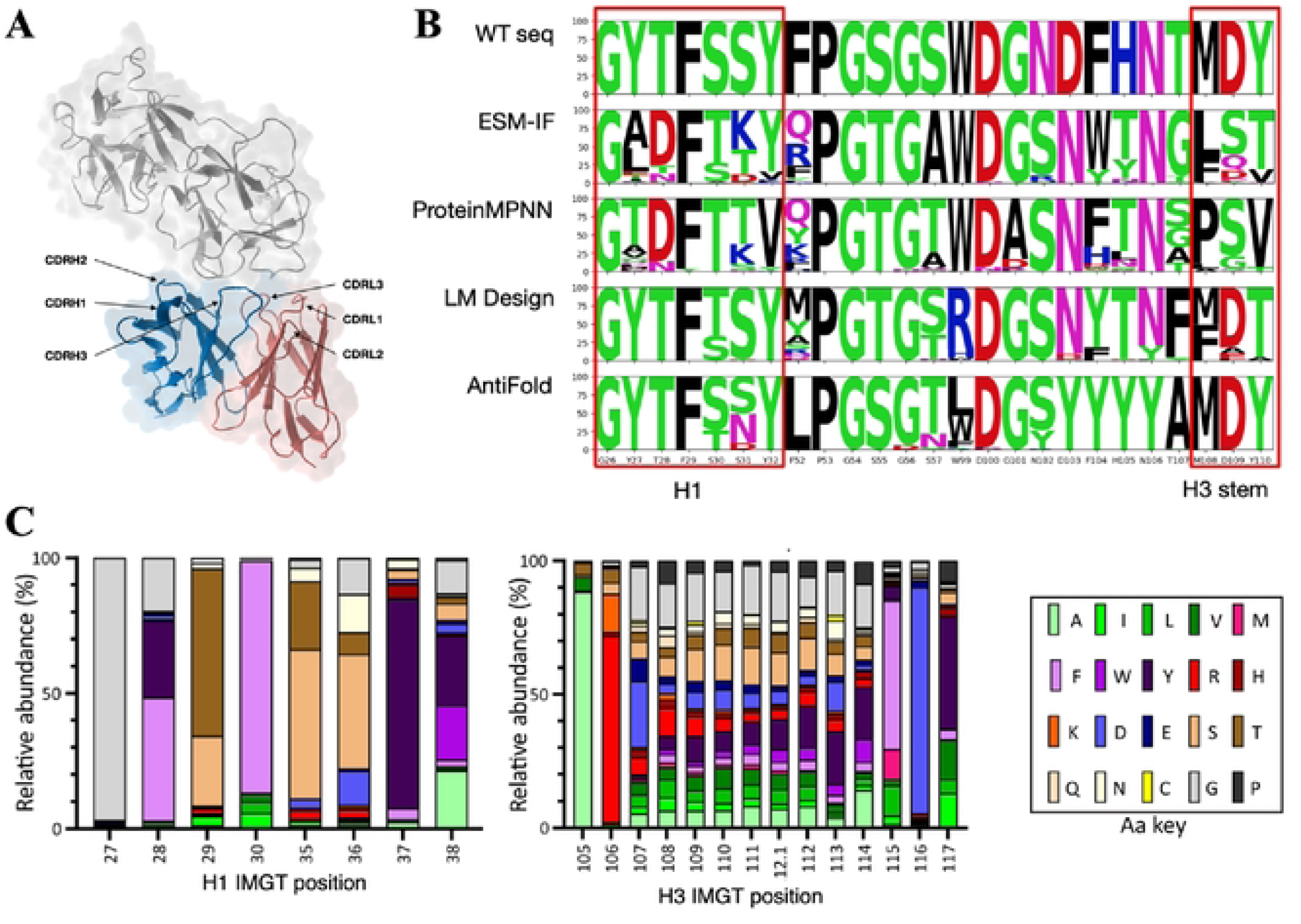
Analysis of CDR Design and Natural Sequence Distributions A) Structural representation of Ricin in complex with Fab SylH3 (PDB ID: 8TFL), highlighting the CDR regions (HI, H2, H3, L1, L2, and L3). B) Sequence logo plots of 100 heavy chain designs generated for PDB 8TFL using different models (ESM-IF, ProteinMPNN, LM Design, and AntiFold). Conserved regions, including H1 and the H3 stem, are highlighted in red boxes. The logo plots illustrate amino acid diversity and conservation at each position. C) Natural amino acid distributions for H1 and H3 positions based on IMGT data from over 11,000 productive human antibody sequences. IMGT numbering is used in the lower panel, which differs from the sequence-based numbering in the upper panel. The color coding corresponds to amino acid types (key provided) The figures were from Steffen Goletz24.

**Figure 8.**
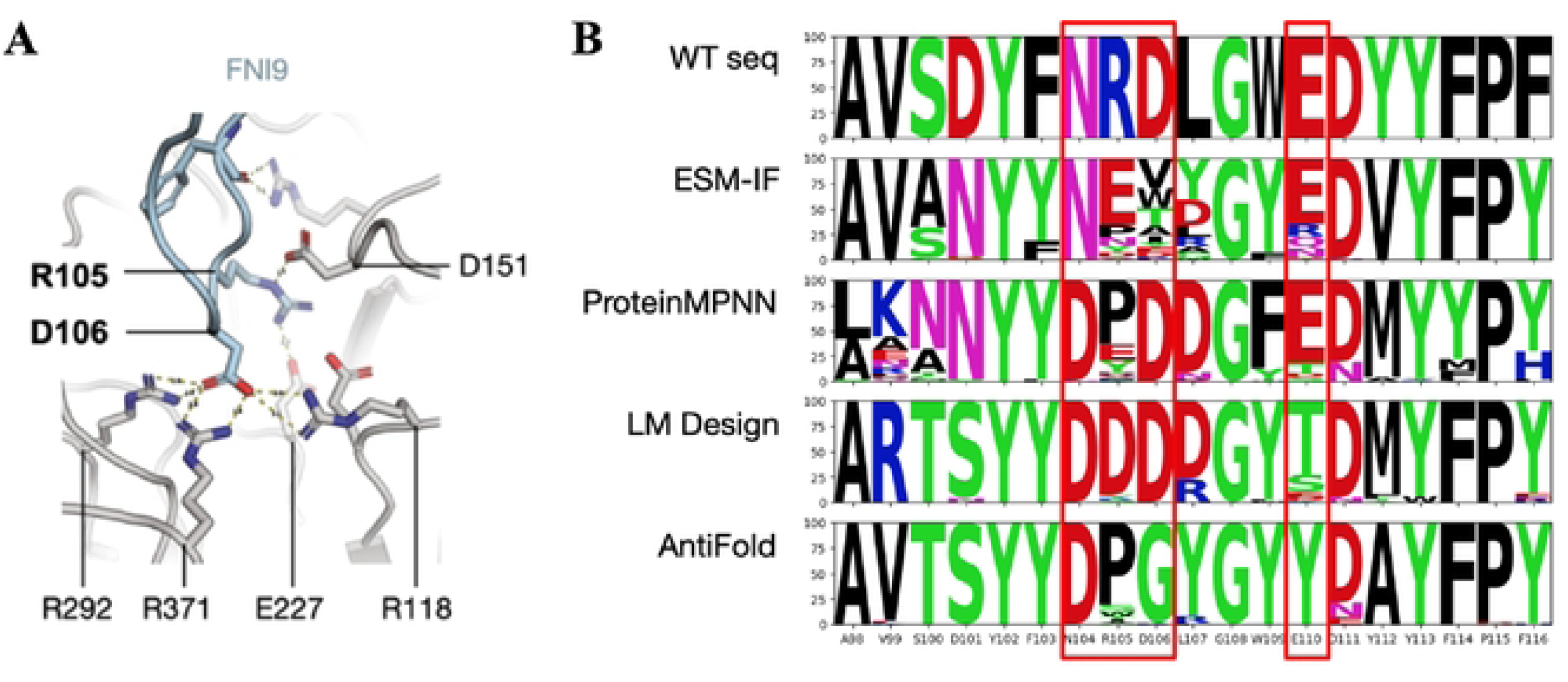
Analysis of Antibody-Neuraminidase Interaction and Heavy Chain Design A) Key interactions between the FNI9 antibody and the neuraminidase catalytic site. The highlighted residues (R105, D106, D151, R292, R371, E227, and R118) indicate critical binding interactions within the epitope. Figure adapted from Matteo Samuele Pizzuto25. B) Sequence logo plots showing the frequency of amino acids in 100 heavy chain designs for PDB 8G3V generated using different models (ESM-IF, ProtcinMPNN, LM Design, and AntiFold). Key epitope-contacting residues arc marked with red boxes, illustrating conservation and substitution patterns across the designs.

For H3 design, AntiFold and LM Design were more successful in accurately designing the stem portion compared to ProteinMPNN and ESM-IF. H1 and the stem of H3 are characterized by relatively conserved sequence patterns, and it is evident that AntiFold and LM Design have effectively learned these conserved patterns for antibody CDRs. For instance, in H1, the G-Y/F-T-F-S/T-S-Y sequence is one of the most frequent patterns, and both LM Design and AntiFold were able to replicate it.

Additionally, we observed that AntiFold achieved a much higher recovery rate for L2 compared to all other methods. For example, in the case of PDB 8TFL, AntiFold reached an 85% recovery rate, while the other methods hovered around 50%. L2 has a limited number of canonical structures, and AntiFold has successfully learned the structure-sequence relationship for this region.

To further evaluate model performance, we investigated instances where models other than AntiFold excelled in H3 design. PDB 8G3V, featuring the FNI9 antibody complexed with neuraminidase, provides a representative example [25]. In this structure, H3 of FNI9 interacts with three highly conserved arginine residues within the neuraminidase catalytic site. Our analysis revealed that AntiFold did not effectively recapitulate these key binding interactions. Specifically, for the critical residues R105, D106, and E110, LM Design and ProteinMPNN successfully recovered D106, while ProteinMPNN and ESM-IF recovered E110. In contrast, AntiFold failed to generate sequences that maintained these interactions, indicating potential limitations in its ability to prioritize functionally critical residues within the H3 loop.

#### Predicting the Effects of Mutations

Beyond sequence design, inverse folding models can also be employed to predict the effects of mutations on antibody binding affinity. By evaluating the log-likelihood of mutant sequences conditioned on the antibody-antigen complex structure, these models can provide insights into the functional consequences of amino acid substitutions.

To benchmark the models’ mutation prediction capabilities, we utilized two types of datasets:

1. SKEMPI2 Database: This database contains a curated set of experimentally determined changes in binding free energy (ΔΔG) for a variety of protein mutations. We chose two antibody-antigen complexes from this database with over 20 mutation data points each [26,27].
2. Deep Mutational Scanning Data: These datasets provide comprehensive mutational landscapes for specific antibodies, typically generated through saturation mutagenesis or combinatorial mutations of CDR regions. The datasets include corresponding binding affinity measurements obtained via high-throughput screening methods. We incorporated several publicly available deep mutational scanning datasets for well-characterized antibodies [17,18].

We used the inverse folding models to compute the log-likelihood scores for each mutation in these datasets, conditioned on the corresponding template PDB structures. We then analyzed the Spearman correlation between the log-likelihood scores and the experimentally measured changes in binding free energy (ΔΔG) or affinity values. We also included two additional language models, ESM2 [28] and xTrimo-1b [29] in the benchmark.

Our analysis revealed that ESM-IF, ProteinMPNN, and LM Design consistently exhibited positive correlations between their log-likelihood scores and the experimentally measured mutation fitness. This suggests that these models can effectively capture the impact of mutations on antibody binding affinity. In contrast, AntiFold and other protein language models showed less consistent correlations, with greater fluctuations across datasets. Notably, ESM-IF demonstrated the highest average correlation across all datasets, indicating its superior performance in predicting the effects of mutations (Figure 9 and Supplementary Figure) This phenomenon likely stems from its design in the training and inference process. The model was trained exclusively on single chains, and for complex inference, it simply concatenates chains. As a result, ESM-IF may misinterpret complex interfaces as analogous to the cores of single-chain proteins during sequence design. This can cause the model to overemphasize mutations that disrupt interface interactions, treating them as though they were internal to a single chain rather than part of a multi-chain assembly.The ESM2 LLM exhibited a remarkable correlation with the CR6261 dataset, which features a diverse array of mutation combinations spanning both the CDR and framework regions. Notably, some of these combinations might negatively impact antibody stability, a factor the LLM seems adept at identifying and capturing [30].

**Figure 9.**
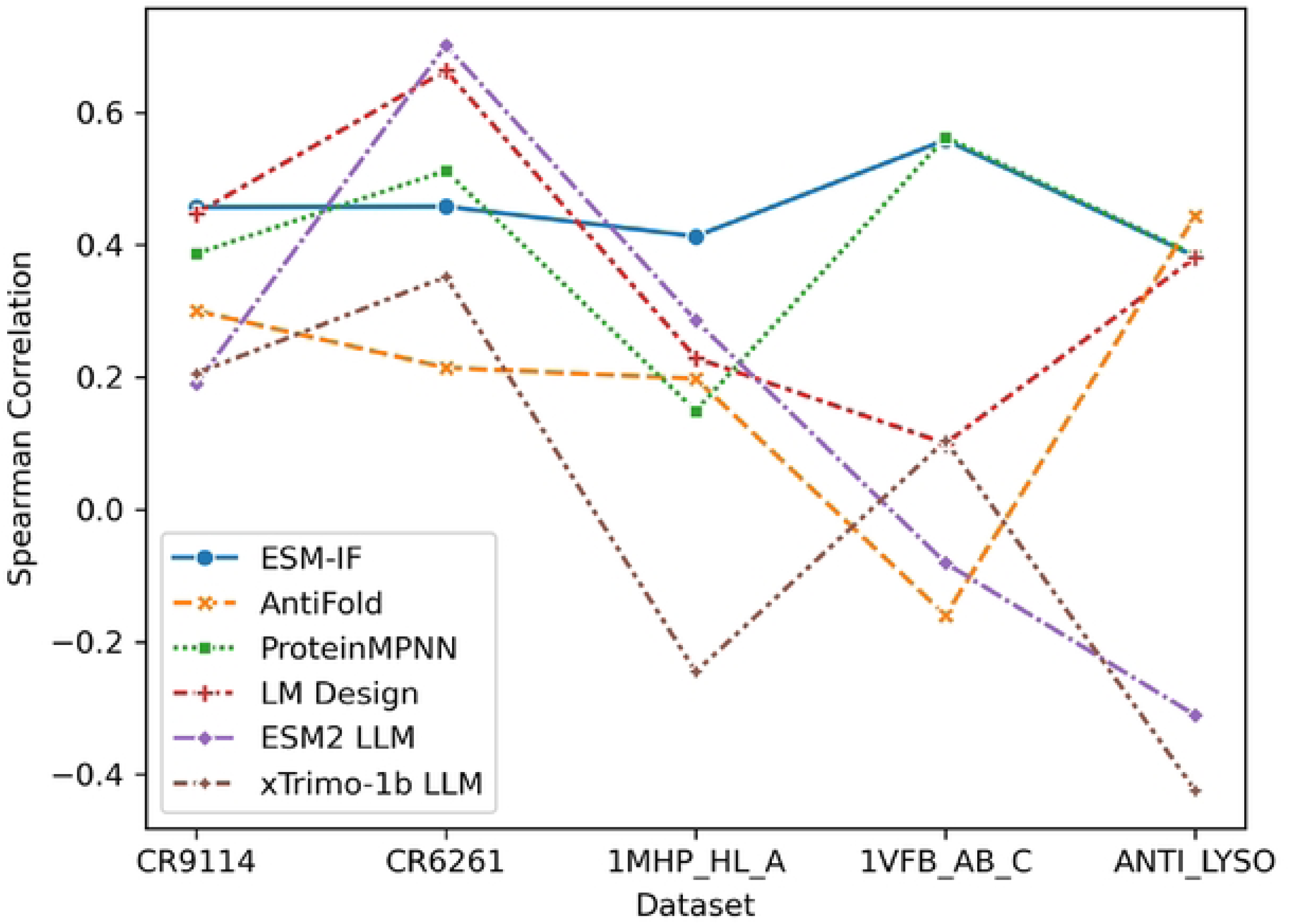
Spearman Correlation Between Model Predictions and Experimental Fitness Scores Spearman correlation coefficients were calculated between the log-likelihood scores predicted by different models and the experimentally measured fitness scores across five independent datasets (CR9114, CR6261, 1MHP_HL_A, 1VFB_AB_C, and ANTI_LYSO). In addition to ESM-IF, AntiFold, ProteinMPNN, and LM Design, two large language models (ESM2 and xTrimo-1b) were included in the benchmark. The results highlight the variability in model performance across datasets and the impact of incorporating advanced language models into the evaluation.

**Figure 10.**
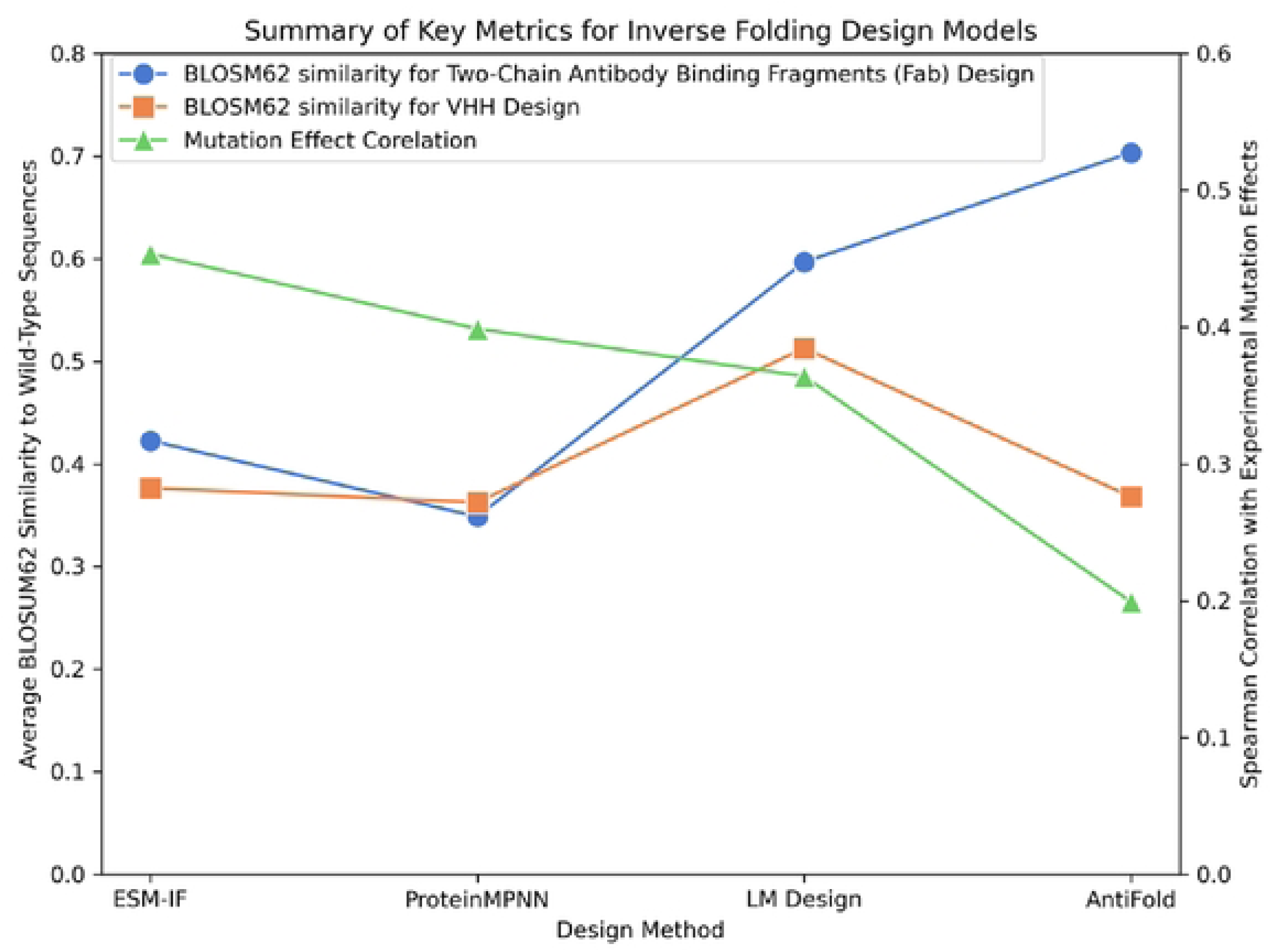
Similarity and Mutation Effect Correlation Across Inverse Folding Design Models This figure summarizes key metrics for inverse folding design models. The BLOSUM62 similarity to wild-type (WT) sequences is shown for Two-Chain antibody binding fragments Fab designs (blue circles) and VHH designs (orange squares) on the left y-axis, while the mutation effect correlation is plotted on the right y-axis (green triangles). All metrics were calculated as the average across all data points in the benchmarking dataset, highlighting model performance across sequence similarity and predictive accuracy for mutational effects.

## Discussion

### Common Features and Learned Properties

Despite performance variations, our benchmark revealed common features among the evaluated inverse folding models. Notably, all models demonstrated an understanding of amino acid physicochemical properties, as evidenced by the confusion matrices. Amino acids with similar properties, such as the aromatic residues F/W/Y, the aliphatic residues I/V/L, and the negatively charged residues D/E, exhibited higher confusion rates within their respective groups. This indicates that the models have learned to associate these residues based on shared characteristics.

Furthermore, all models consistently achieved high recovery rates for proline and glycine. LM Design, in particular, excelled in predicting these residues, likely due to their unique dihedral angle distributions. This suggests that the models leverage structural information encoded in these distinct dihedral angle preferences during sequence design.

### Distinct Model Characteristics and Training Data Influences

#### Antibody-Specific Features

The performance of inverse folding models is heavily influenced by the specificity of their training datasets. AntiFold and LM-Design consistently demonstrated higher sequence recovery rates for antibody-specific features, particularly in conserved regions such as H1 and the C-terminus of H3. This suggests these models effectively learned antibody-specific structural and sequence motifs during training. For example, AntiFold’s fine-tuning on antibody-specific datasets, such as SAbDab and modeled structures from OAS, enabled it to capture the unique sequence-structure relationships present in antibody frameworks.

Conversely, LM-Design benefits from its integration of a protein language model, which likely helps refine its antibody-specific predictions during inference. Despite being trained on general protein datasets, the additional structural encoder enables LM-Design to incorporate antibody-specific features effectively.

#### Limitations of General Protein Data

Models such as ProteinMPNN and ESM-IF, trained on general protein datasets, showed limitations in capturing antibody-specific nuances. For example, these models often struggled to generate sequences for H3, which is highly variable and critical for antigen binding. This limitation likely arises from their reliance on datasets dominated by non-antibody proteins, where such structural diversity is less prevalent.

Furthermore, general protein training data may lead to biases in sequence design. For instance, ProteinMPNN and ESM-IF exhibited overrepresentation of high-frequency amino acids (e.g., serine, glycine) that are common across general proteins but less contextually appropriate for certain antibody regions.

The differences in model performance highlight the need for curated training datasets to improve antibody-specific design. AntiFold’s poor performance on VHH sequences underscores this issue. As VHH antibodies were largely absent from its training set, the model struggled to generate sequences for these structures, demonstrating reduced generalizability across antibody subclasses.

To address such limitations, future models should incorporate broader datasets that include VHH antibodies and antigen-antibody complexes. Additionally, integrating functional data such as binding affinities and mutational landscapes could enhance models’ ability to prioritize residues critical for binding, particularly in H3 regions.

Finally, the reliance on antigen information also varies among models. AntiFold, for instance, displayed minimal dependence on antigen structure, focusing primarily on the CDR loop itself. While this improves its robustness in antigen-independent tasks, it may reduce its utility for designing antigen-specific interactions, a critical consideration for therapeutic applications.

#### Future Directions for Antibody Inverse Folding

While antibody inverse folding models have demonstrated promising progress, there remains significant potential for improvement. Below, we expand on five key areas for advancing this field, with practical examples and potential impacts highlighted.

**Enhanced Training Data:** Models trained on general protein data may not fully capture antibody-specific nuances, particularly in H3 design as shown in this study. Curating larger, more diverse antibody-specific datasets, including VHH sequences and broader antigen-antibody complexes, could improve model performance and generalizability.

**Integration of Functional Data:** Incorporating functional data, such as binding affinities and mutational effects, could enhance model accuracy and informativeness e.g. the model could focus more antibody-antigen interactions when designing the interface residues. This could be achieved through multi-task learning or by developing models that explicitly consider both structural and functional constraints.

**Improved Generalization:** Addressing limitations in generalizing to different antibody types and structural contexts is crucial. Techniques such as transfer learning, data augmentation, and incorporating diverse structural templates during training could improve model generalization.

**Comprehensive Evaluation Metrics:** Developing more comprehensive evaluation metrics that capture the functional and biophysical properties of designed antibodies is essential. This could involve incorporating experimental validation, biophysical simulations, and machine learning-based predictors of antibody properties.

**Mitigation of Biases:** Addressing potential biases in the models, such as overrepresentation of certain amino acids, is necessary. Adjusting training data distributions, incorporating regularization methods, or developing models that explicitly account for amino acid usage biases could mitigate these biases.

By addressing these areas, future research can drive the development of more accurate, generalizable, and robust inverse folding models, facilitating the design of novel antibodies with desired properties for therapeutic and biotechnological applications. Generative LLMs for antibody design have the potential to accelerate drug discovery and design by delivering better and faster lead candidates e.g., desired epitope, no liabilities that could be bespoke to individual patients needs and thereby enable next generation of therapeutics.

## Contribution

YLi conducted the majority of sequence design, data analyses, and drafted the manuscript. YL performed sequence design using ProteinMPNN, AbMPNN, and ESM-IF, and executed the relaxation protocol. CX contributed to the development of the RDE mutation effect prediction pipeline and participated in the results discussion. YZ assisted in curating the benchmarking dataset and contributed to the results discussion. ZP supported the results discussion. PJG supervised the project, analyzed model performance, and drafted the manuscript.

## Acknowledgement

The authors would like to thank Dr. Matthew Raybould, Yuan Wei, and Patrick Carney for nice comments and suggestions. The authors would like to thank Bing Yang for helpful discussion of the model training strategy.

